# Object Speed Perception during Self-Motion in Depth

**DOI:** 10.64898/2026.06.20.733496

**Authors:** Anita Pandey, Ahmed Nadeem, Laurence R. Harris, Björn Jörges

## Abstract

During sideways movement of an observer, optic flow parsing – in which an object’s speed in the world is extracted from all the other visual movement present in the scene, self-generated and otherwise – has been shown to be incomplete, leading to biases in speed perception, particularly when object and observer are moving in opposite directions. Here, we assess how judgements about the speed of objects moving in depth (judged relative to the world) towards or away from an observer (6 m/s) are affected by simultaneous movement of the observer either in the same or opposite direction as the object. In a virtual reality display, participants (n = 25) viewed a sphere simulated as moving in a corridor either while they were stationary or during visually simulated self-motion in the same or opposite direction as the object. They judged the sphere’s movement relative to the world by comparing its motion to a probe sphere that travelled laterally across the corridor in front of them. In a second experiment (n = 28) participants performed the same task but during faster self-motion (10 m/s). The second cohort also judged the direction in which the object was perceived to move during the same combinations of self and object speeds. Object speed was overestimated when the object travelled in the direction opposite to the observer compared to how object’s motion was judged when the observer was stationary. However, object speed was also overestimated during self-motion in the same direction as the object where participants were also much more likely to misjudge the direction of motion of the object. Precision of judgements was lower when self-motion was simulated than it was for stationary observers. A simple arithmetic model of flow parsing fails to capture these results satisfactorily, suggesting that different mechanisms may be at play when the observer travels in the same direction as a moving object and is vulnerable to misperceiving its direction of travel.

## Introduction

How accurately are we able to estimate the speed of a moving object when we ourselves are in motion? The process of gauging object motion during self-motion requires us to subtract or cancel out the retinal motion created by our own self-motion from the motion of the moving object. This subtraction is referred to as *flow parsing*. According to the Flow Parsing Hypothesis (Rushton and Warren 2005; Warren and Rushton 2008, 2009b; Dupin and Wexler 2013; Rushton et al. 2018) the observer’s task is to determine the optic flow created by their own motion and then subtract this from the overall retinal input. This would then leave the impression of a static environment in which any remaining motion can be attributed to object motion. This model corresponds well to our everyday experience of moving around the world while watching dogs, balls, people and cars moving around. The Flow Parsing hypothesis has been demonstrated to accurately model the perception of object motion during self-motion – with some limitations – for up/down translation (Dyde and Harris 2008), rotation (Probst et al., 1995; Garzorz et al., 2018) and for motion-in-depth (Aguado & Lopez-Moliner, 2019; Dupin & Wexler, 2013; Gray et al., 2004).

During visually simulated self-motion in the laboratory, in the absence of actual physical motion, the self-motion component tends to be underestimated (Dokka et al. 2015). This may be due to a dissonance between the visual and vestibular systems since the participant does not actually move and their vestibular system is consequently quiet (Becker et al. 2002; Frissen et al. 2011). Any underestimation of self-motion when the observer’s motion is visually simulated should then lead to a bias in the estimation of object-motion seen during such simulated self-motion because the observer would attribute less of their overall retinal input to their own motion leading to incomplete flow parsing and an overall bias in estimating the speed of object motion relative to the world. And in fact, flow parsing has been shown to be incomplete in a large number of studies (Rushton and Warren 2005; Warren and Rushton 2008, 2009b, a; Jörges and Harris 2022, 2024). Furthermore, parsing out the consequences of self-motion and object motion from optic flow tends to decrease precision (Dokka et al. 2015; Jörges and Harris 2024).

Older studies on flow parsing have tended to focus on time-to-contact estimation during self-motion or direction judgements or motion detection (Rushton and Warren 2005; Warren and Rushton 2008, 2009a). Largely absent from this literature are direct assessments of perceived *speed* during self-motion (c.f., Hogendoorn, Alais et al., 2017). We have previously shown that speed judgements of laterally moving objects seen during lateral self-motion are biased according to the predictions of the flow parsing hypothesis (Jörges and Harris 2022) but only when object and observer move in opposite directions. For same-direction motion, no biases in speed judgements were found. In a follow-up study (Jörges and Harris 2024), we replicated this finding but showed that, in a time-to-contact estimation task, the same participants displayed flow parsing biases both for opposite direction *and* same direction motion profiles. This dissociation between speed judgements and the more action-oriented time-to-contact judgements highlights how the effectiveness of flow parsing can depend on the task. While flow parsing in depth has been studied thoroughly using time-to-contact estimation and direction estimation tasks, assessing the effectiveness of flow parsing when judging the speed of objects approaching or moving away from an observer has not been systematically investigated.

Here, we assess the effectiveness of flow parsing in depth during visually simulated self-motion using a speed estimation task. Given that, due to the absence of physical motion, self-motion is expected to be underestimated, we expect flow parsing to be incomplete, leading to an overestimation of object speed relative to the world when observer and target move in opposite directions and to an underestimation of object speed when observer and target move in the same direction. Due to the added computations required to extract object motion from retinal flow during observer motion, we further expect precision to be lower during self-motion than for stationary observers.

## Methods

### Participants

We recruited 25 participants for Experiment 1 and 28 participants for Experiment 2, for a total of 53 participants through the Undergraduate Research Participant Pool (URPP) at York University, Toronto, Canada. Participants received course credit as compensation for their participation. There were no exclusion criteria for participating in this experiment and this study received ethics approval from the Human Participant Ethics Review Sub-Committee at York University. Informed consent was obtained from all the participants in writing, and the experiment was conducted in accordance with the principles of the Declaration of Helsinki.

### Apparatus

The experiment was performed in virtual reality with participants remaining physically static and seated. The stimulus was programmed in Unity (2019.2.11f1) and both the Unity projects and the executables for both experiments can be found on Open Science Foundation (https://osf.io/ty7cg/). A tower PC with an Intel® Core™ i7-8700 CPU @ 3.70GHz, 16GB, 2666 MHz RAM, and an 8GB NVIDIA GeForce RTX 2070 graphics card was used. The stimulus was presented stereoscopically using an HTC VIVE Pro Eye VR headset and participants responded with a keyboard. We used R v4.4.2 for graphics, statistical analyses and modelling.

### Stimulus

The VR environment was presented in stereo and included depth cues from the light source, shadows, as well as from the range of textures on the floor and walls. The environment consisted of tiled walls to the left and right to the observers and small potted plants distributed across the tiled floor, as seen in Figure 1.

**Figure 1:**
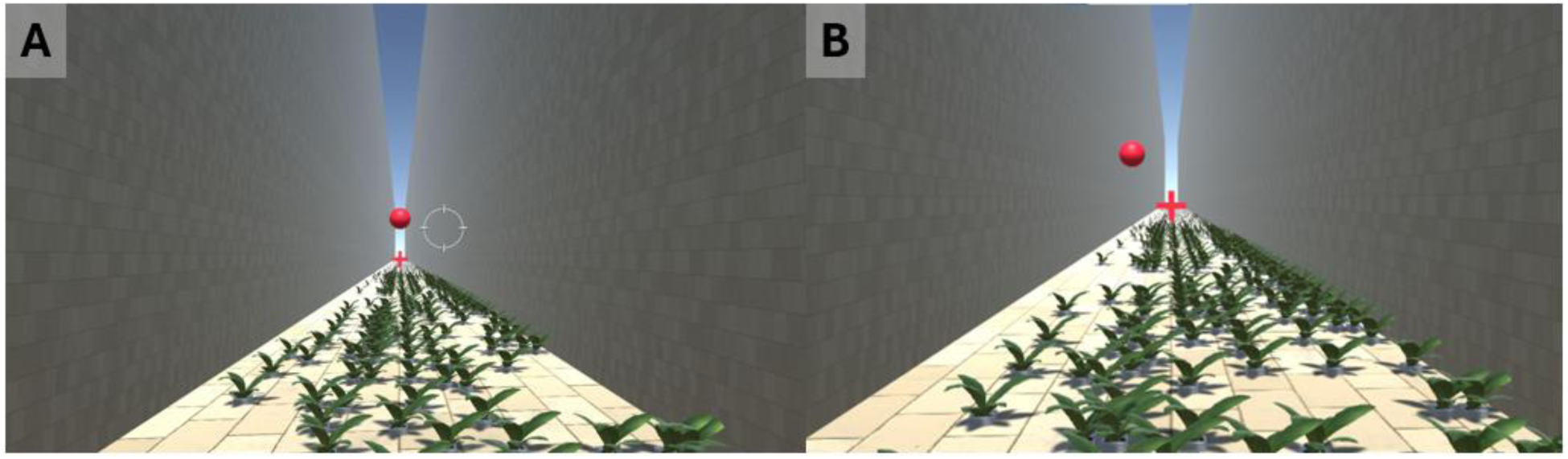
Screenshot of the experiment environment in virtual reality (VR) as seen through the headset. (A) shows the target sphere that moves either toward or away from the observer. (B) Shows the probe sphere that moves laterally across the corridor. The tiled pattern on the walls is barely visible in these photos but can be clearly seen in the video from the 6 m/s self-motion experiment which is available on the Open Science Foundation repository: https://osf.io/ty7cg/

### Object Motion

While immersed in this environment, participants were asked to judge the speed of a target sphere that moved either towards or away from them (Fig 1a) relative to a probe sphere that moved laterally across the corridor (Fig 1b). Was the probe ball faster or slower? In order to do this, they needed to estimate the speed of the target sphere relative to the world. Both spheres were red, had diameters of 40 cm and moved at a height of 2 m above the simulated ground, placing them above the observer’s simulated eye height and reducing the impression of a direct collision with the observer. The target sphere moved at either 3 m/s or 10 m/s, either towards the observer or away from them. The starting point was generated such that the closest distance between the observer and the sphere was 0.5 m, which could either occur at the beginning or at the end of presentation. The probe sphere moved either from left-to-right or from right-to-left (randomly chosen on each trial) across the 8m-wide corridor in front of the observer at a speed that was governed by two interleaved PEST staircases (Taylor and Creelman 1967) one of which started at a speed 30% above the target sphere’s speed, while the other started at 30% below the target sphere’s speed. Each PEST was terminated after 15 trials.

### Self-Motion

The participants’ viewpoint was simulated at 1.3 m above the ground (i.e., 70 cm below the path of the target and probe spheres), which corresponds to an average adult’s seated eye-height. There were three conditions of observer motion: either stationary, moving backwards, or moving forwards at either 6 m/s (Experiment 1) or 10 m/s (Experiment 2). All self-motion was passive and visually simulated.

### Head Angle Control

Eye tracking was not included in this experiment, but we implemented a head direction control to ensure that participants were consistently facing forwards down the corridor. In the beginning of each trial participants aligned a white target circle with the red fixation cross by adjusting the position of their head so that the target circle aligned with the cross (see Figure 1). Once aligned with the cross (i.e., within 2.5° of the target), the target circle disappeared but reappeared if the participant’s head drifted. After starting any given trial, head deviations did not lead to trials being ended or restarted as participants were not assumed to make major head movements throughout the short period of the trial.

### Procedure

Participants were seated in front of a laptop in the lab setting and fitted with the VR headset. After receiving general instructions on the task, they then performed a short test to make sure they had understood the controls: they were shown one trial in which the target sphere moved towards or away from them much faster than the probe sphere and one trial in which the target sphere was moving much slower than the probe sphere. Only after they had answered correctly on both trials, they were allowed to proceed to the main experiment. They then completed the main experiment, which interleaved trials pertaining to all conditions (three simulated self-motion conditions: forwards/backwards/stationary; two sphere speeds: 3 m/s and 10 m/s; two sphere directions: towards the observer/away from the observer). With two staircases per condition and 15 trials per staircase, participants completed a total of 3 x 2 x 2 x 2 x 15 = 360 trials. Staircases were regular PESTs (Taylor and Creelman 1967). The experiment took around 30 to 40 minutes to complete, including setup, instructions and training. Participants were allowed to take breaks at their own discretion.

### Experiment 1 and Experiment 2

We collected data in two very similar versions of this experiment that differed only in the simulated speed of self-motion. The participants of Experiment 1 were simulated to be moving at 6 m/s, while the participants in Experiment 2 were moving at 10 m/s.

### Perceived Object Direction

For Experiment 2, we also added a short secondary experiment in which we asked participants whether they thought the target sphere was moving in the same direction as they were or in the opposite direction. i.e., we asked them to judge the sphere’s movement relative to the world. We showed them each combination of self-motion speed (0 m/s, 10 m/s forwards, 10 m/s backwards) and sphere speed (3 m/s towards the observer, 3 m/s away from the observer, 10 m/s towards the observer, 10 m/s away from the observer) 10 times (for a total of 3 x 4 x10 = 120 trials) and asked them after each trial to indicate whether they thought the sphere was moving in the same direction as they were or in the opposite direction. This task was always executed after the speed judgement tasks and took about 5 minutes to complete.

## Data Analysis

### Fitting Psychometric Functions

We used the R package quickpsy (Linares and López-Moliner 2016) to fit cumulative normal functions to all the data from the two interleaved PESTs for each condition using a direct likelihood maximization approach. This allowed us to use all the data for a given participant and condition and gave us a reliable estimate of precision. The mean of the fitted cumulative normal functions corresponds to the Point of Subjective Equality (PSE), i.e., the stimulus velocity at which participants could not tell whether the probe or stimulus object was faster, and the fitted standard deviation corresponds to the 84.6% Just Noticeable Difference (JND), as a measure of precision. Fitting the function allowed us to use all the data from the two staircases and also provided the precision estimate.

### Outlier Analysis

We conducted an outlier analysis over the fitted PSEs and JNDs. We first excluded impossible values, i.e. PSEs below 0 and JNDs of 0 or lower, as the result of bad fits. This criterion led to the exclusion of 118 out of 636 total conditions (18.6%) across both experiments. We then also removed all conditions where the fitted PSE or JND were 1.5 times the interquartile range below/above the 1^st^/3^rd^ quartile of the respective distributions across all participants and conditions (82 of the remaining 518 conditions, or 15.8%, across both experiments), for a total of 436 (or 68.6%) included conditions.

### Statistical Analysis

We then used Linear Mixed Modelling (as implemented in the lme4 package (Bates et al. 2015) for R (R Core Team 2017)) to test our hypotheses regarding biases and precision differences in response to self-motion. To test for biases, we fitted a model with the PSEs as dependent variable, the sphere speeds (3 m/s and 10 m/s), the self-motion condition (same direction, fast; same direction, slow; opposite direction, fast; opposite direction, slow; no self-motion) as well as their interaction as fixed effects and random intercepts per participant as random effects. In Wilkinson & Rogers (1973) notation, this reads as follows:

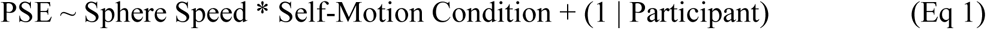

We fitted the same model to assess precision differences, but with the JNDs as dependent variables:

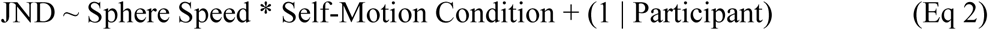

We then computed 95% confidence intervals to determine statistical significance using the confint function from base R.

To test for differences in perceived direction, we included trials pertaining to conditions that were also included into the main PSE/JND analysis. We then employed generalized linear mixed modelling with a logit link function to determine condition-wise differences in how likely participants were to identify the direction of the sphere correctly. We used the participants’ responses (Correct/False) as dependent variable instead of PSEs/JNDs and employed the same random effects structure as for the PSE/JND analysis:

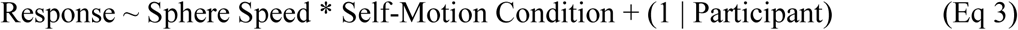

As for the other analyses, we computed 95% confidence intervals to assess statistical significance.

## Results

### PSEs

Figure 2 shows the PSEs (perceived speeds relative to the world) for all conditions in both experiments and Table 1 provides the full difference contrasts and 95% confidence intervals for both PSE linear mixed models. Note that all values are positive as we only asked if the sphere’s motion was faster or slower than the probe sphere, not their perceived direction. Overall, the faster target speed (10m/s) evoked faster PSEs in both experiments and in both directions. Notably, the PSE difference between sphere speeds of 3 m/s and 10 m/s was only about 2.2 m/s for Experiment 1 and 1.6 m/s for Experiment 2 for stationary observers: much lower than the actual difference between the target speeds. Curiously, all self-motion conditions were associated with significantly faster PSEs compared to when observers were stationary. This was predicted for the Opposite Directions conditions (where it corresponds to incomplete flow parsing) but was highly unexpected for the same direction conditions where the relative speed between target and observer would have been *lowered* by the simultaneous self-motion.

**Figure 2:**
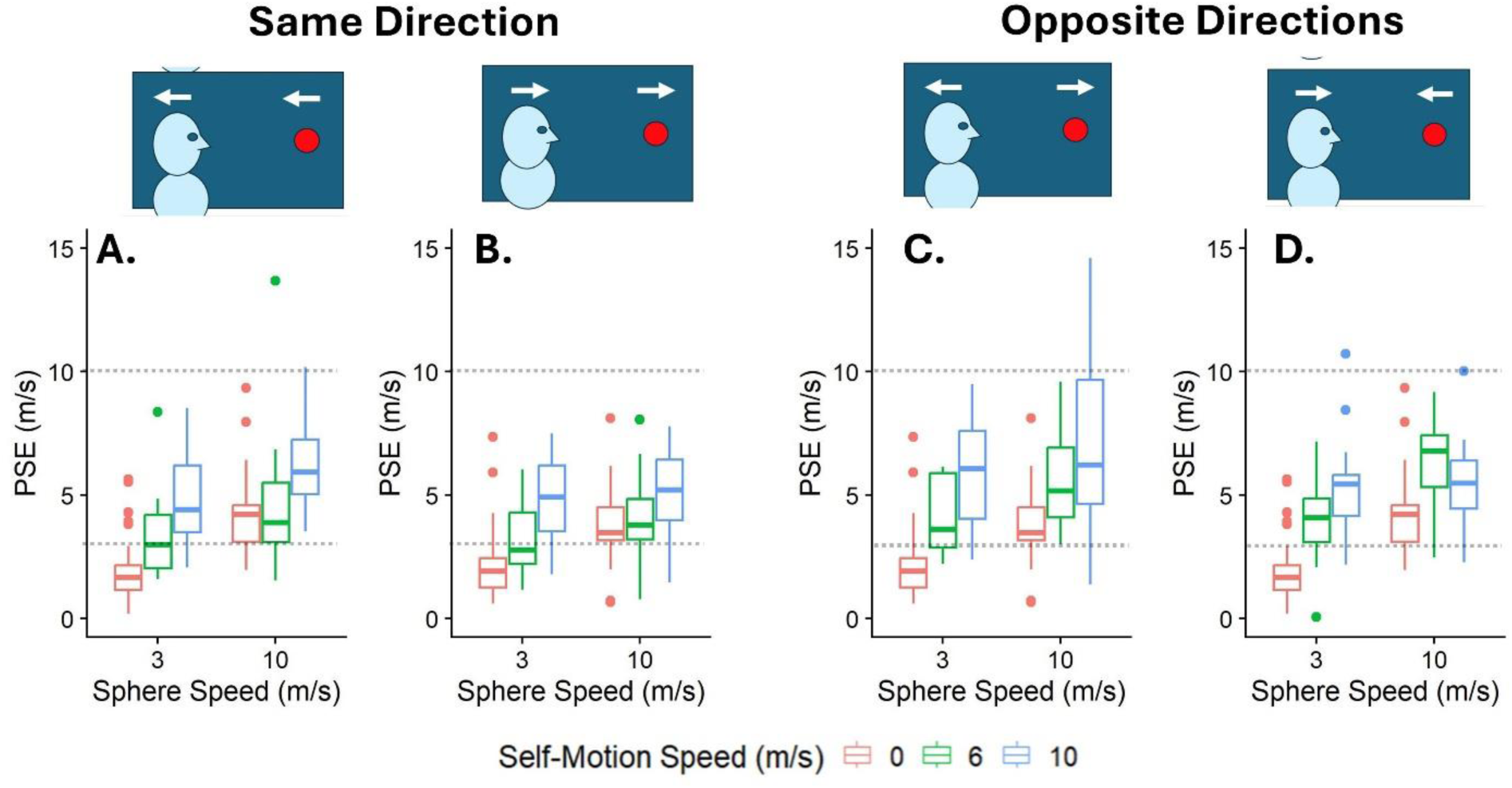
Boxplots of the PSEs for sphere speeds (x axis), separately for each combination of observer and sphere speed (panels) and simulated self-motion speeds (color-coded). A. For observer and sphere motion out of the scene, i.e., in the direction the participant’s back was facing, as shown by the arrows in the cartoons. B. For observer and sphere motion into the scene, i.e., down the corridor as shown by the arrows. C. For backward observer motion and sphere motion down the corridor, i.e., in the same direction as the observer was facing as shown by the arrows. D. For forwards observer motion and sphere motion up the corridor. The dotted horizontal lines at 3 and 10m/s correspond to the actual speeds at which the balls were moving.

**Table 1:** Differences contrasts and 95% confidence intervals for all statistical comparisons in the speed judgement tasks. Significant positive effects are color-coded in bold black and significant negative effects are color-coded in bold red.

|  | Difference Contrast (m/s) | 95% CI Lower (m/s) | 95% CI Upper (m/s) | Signif. |
| --- | --- | --- | --- | --- |
| ***PSEs – Experiment 1 (6 m/s Self-Motion)*** |  |  |  |  |
| Sphere Speed 10 m/s (versus 3 m/s) | **2.22** | **1.76** | **2.7** | * |
| Opposite Direction (versus Stationary) | **2.17** | **1.71** | **2.67** | * |
| Same Directions (versus Stationary) | **1.42** | **0.96** | **1.87** | * |
| Sphere Speed * Opposite Directions | -0.09 | -0.85 | 0.59 | n.s. |
| Sphere Speed * Same Direction | **-1.23** | **-1.87** | **-0.54** | * |
| ***PSEs – Experiment 2 (10 m/s Self-Motion)*** |  |  |  |  |
| Sphere Speed 10 m/s (versus 3 m/s) | **1.6** | **1.04** | **2.2** | * |
| Opposite Direction (versus Stationary) | **3.18** | **2.68** | **3.73** | * |
| Same Directions (versus Stationary) | **2.68** | **2.02** | **3.38** | * |
| Sphere Speed * Opposite Directions | **-0.84** | **-1.61** | **-0.04** | * |
| Sphere Speed * Same Direction | **-1.01** | **-1.83** | **-0.13** | * |
| ***JNDs – Experiment 1 (6 m/s Self-Motion)*** |  |  |  |  |
| Sphere Speed 10 m/s (versus 3 m/s) | 0.3 | -0.18 | 0.78 | n.s. |
| Opposite Direction (versus Stationary) | 0.46 | -0.01 | 0.94 | n.s. |
| Same Directions (versus Stationary) | **0.65** | **0.16** | **1.09** | * |
| Sphere Speed * Opposite Directions | 0.61 | -0.06 | 1.26 | n.s. |
| Sphere Speed * Same Direction | -0.66 | -1.34 | 0.06 | n.s. |
| ***JNDs – Experiment 2 (10 m/s Self-Motion)*** |  |  |  |  |
| Sphere Speed 10 m/s (versus 3 m/s) | 0.43 | -0.22 | 1.08 | n.s. |
| Opposite Direction (versus Stationary) | **0.99** | **0.34** | **1.64** | * |
| Same Directions (versus Stationary) | **1.17** | **0.55** | **1.84** | * |
| Sphere Speed * Opposite Directions | 0.12 | -0.91 | 1.01 | n.s. |
| Sphere Speed * Same Direction | -0.91 | -1.89 | 0.04 | n.s. |

The interactions show that the difference in PSEs between the sphere speeds decreased significantly in the presence of self-motion (compared to a stationary observer) for three out of four self-motion conditions (Experiment 1, in Opposite Direction had no significant difference).

### JNDs

Figure 3 shows the JNDs for all conditions in both experiments and Table 1 shows all the difference contrasts and 95% confidence intervals relating to this analysis. Faster sphere speeds were not related to significantly higher JNDs, but simulated self-motion led to higher JNDs in three out of four comparisons. No evidence was found for an interaction between sphere speed and self-motion condition.

**Figure 3:**
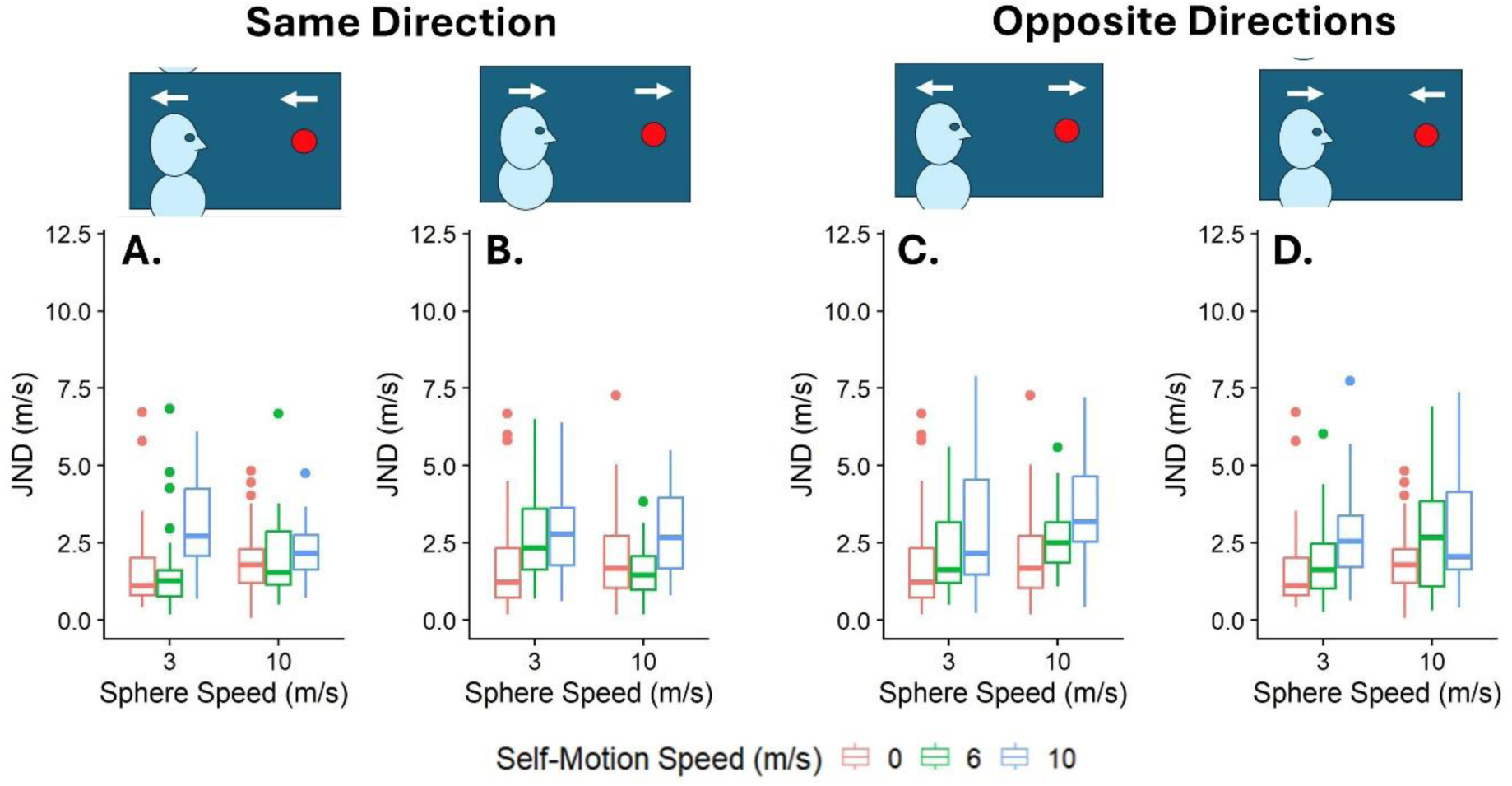
Boxplots of the JNDs for sphere speeds (x axis), separately for the combinations of observer and sphere speeds (panels) and the self-motion speeds (color-coded). Convention as for Fig 2.

### Directions (Experiment 2)

Figure 4 shows the distribution of responses in the direction judgement task. Participants were significantly less likely to give correct responses in the “Same Direction” conditions and significantly more likely in the “Opposite Directions” condition. Both effects were significantly weaker for the higher sphere speed. See Table 2 for the detailed breakdown of all tested difference contrasts as well as the 95% confidence intervals. Participants judged the direction of the sphere incorrectly in more than 80% of the cases when the sphere was travelling at 3m/s in the same direction as the observer.

**Figure 4:**
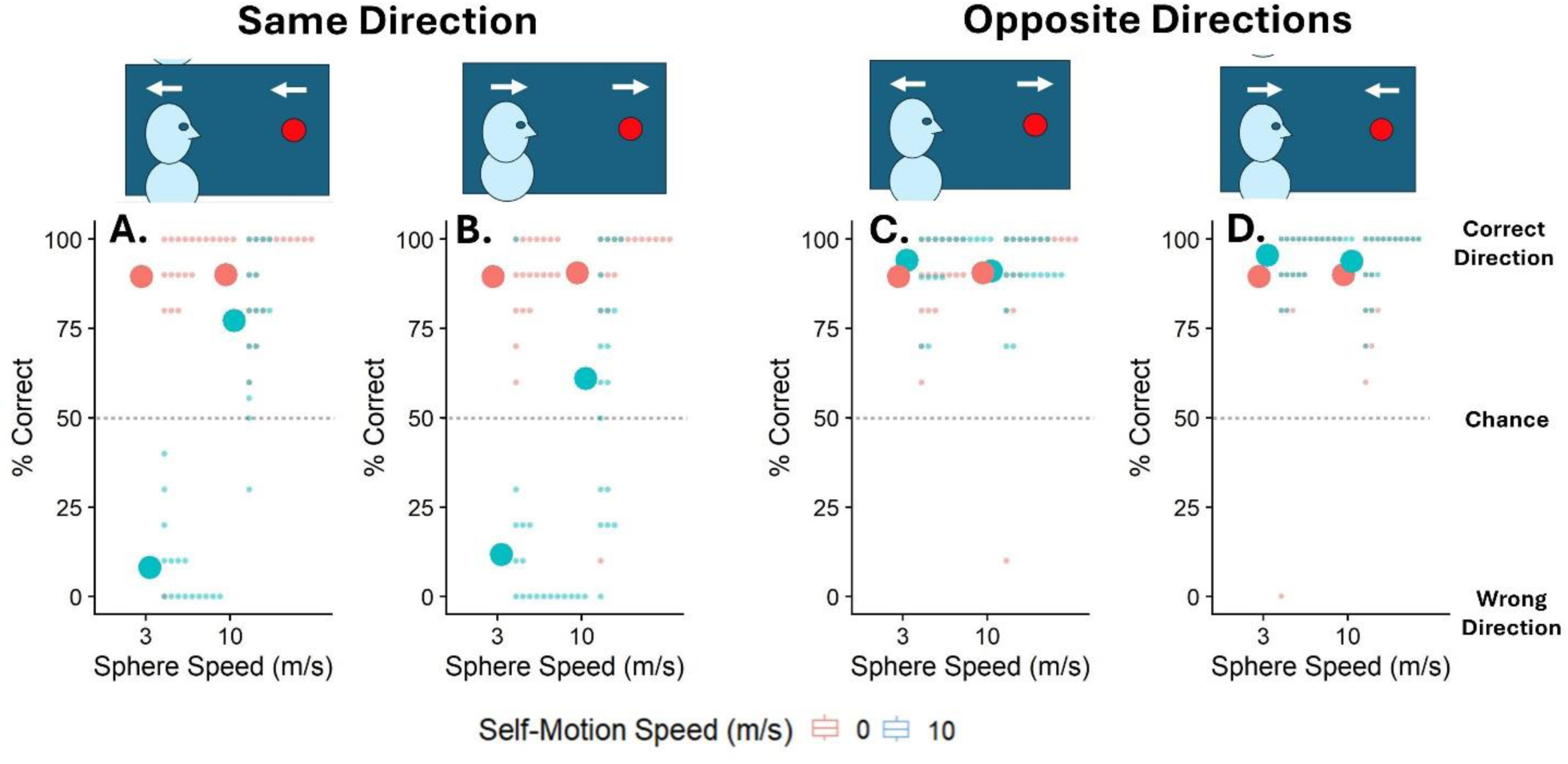
Probability for each participant (distributions of small dots) and mean probability across the whole sample (large dots) for giving the correct response for the direction-judgement task (y axis), split up by sphere speed (x axis), simulated self-motion speed (color-coded). Panels correspond to the relationship between observer direction and sphere direction as in Figures 2 and 3.

**Table 2:** Differences contrasts and 95% confidence intervals for all statistical comparisons in the direction judgement task. Significant positive effects are color-coded in bold black and significant negative effects are color-coded in bold red.

|  | Difference Contrast (odds ratio) | 95% CI Lower (odds ratio) | 95% CI Upper (odds ratio) | Signif. |
| --- | --- | --- | --- | --- |
| **Direction Judgements (Experiment 2)** |  |  |  |  |
| Sphere Speed 10 m/s (versus 3 m/s) | 0.33 | -0.11 | 0.85 | n.s. |
| **Opposite Direction (vs Stationary)** | **0.89** | **0.4** | **1.52** | * |
| **Same Directions (vs Stationary)** | **-4.84** | **-5.52** | **-4.33** | * |
| **Sphere Speed \* Opposite Directions** | **-0.91** | **-1.72** | **-0.23** | * |
| **Sphere Speed \* Same Direction** | **2.93** | **2.2** | **3.69** | * |

### Modelling

We fit a flow parsing model to investigate whether our data supports simple flow parsing. We restricted this analysis to the data collected in Experiment 2 where we had access to independent direction estimates for each participant. We used this formula:

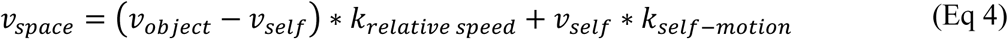

where *v_space_* denotes the perceived object speed in space, *v_object_* represents the actual object speed in space, *v_self_* denotes the actual simulated self-motion speed in space. *v_object_* − *v_self_* is the difference between the object and observer speeds, i.e., the object’s speed on the retina.

*k_relative speed_* then denotes the gain of the internal representation of the speed of the object relative to the observer. This estimate was then added to the perceived speed of the person given by *v_self_* multiplied by the gain with which this is processed: *k_self_*_−*motion*_. We used data from the directions task from Experiment 2 to assigned perceived directions to the PSEs, such that positive values meant that participants perceived the sphere to be moving into the scene and negative values meant that they were perceiving the sphere to be moving out of the scene, i.e., in the direction their back was facing. We assigned the correct direction to all speed judgement conditions except for the condition where the observer moved in the same direction as the object and the object moved at 3m/s which was the condition associated with a reversal of perceived direction (see Fig 4). The relative speed gain (*k_relative_* _*speed*_) was computed – for each participant individually – as the mean PSE in the stationary condition divided by the presented sphere speed:

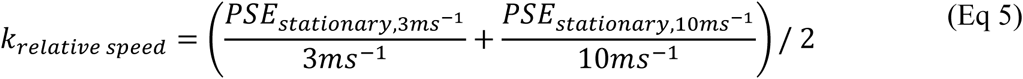

*k_relative speed_* describes the conversion of the relative motion between target and observer into world-relative object speed. This can then be used when evaluating Eq 4.

The median fitted relative speed gain was low with a value of *k_relative speed_* = 0.43, i.e., the sphere moving in depth was perceived at 43% of the speed of the laterally moving probe sphere by a typical participant.

We used this base model (Eq 4) to address several questions:

- Overarching Question: How well can this simple flow parsing model account for the paradoxical finding that object speed was overestimated for the Same Direction motion profile?
- Supplemental Question 1: Do participants flow-parse equally between the Same Direction and the Opposite Directions motion profiles?
- Supplemental Question 2: Is the same perceptual mechanism at work when participants estimate the object direction correctly compared to when they misperceive it (i.e., in the Same Direction, 3m/s condition)?

To address these questions, we fitted the following model variations:

### Fitting the Model (One-parameter fits)

We fitted the model by minimizing the mean squared error (MSE) between predicted and (signed) observed data separately for each participant. *v_object_* and *v_self_* were given by the experimental design, *k_relative speed_* was computed for each participant separately as per Equation 5 and the remaining one free parameter, *k_flow_*_−*parsing*_ was then fitted to the data. We used the optim function from base R with the built-in BFGS optimization method. We restricted the gains to the interval [-1; 2]. While values below 0 and above 1 are theoretically dubious, inherent noise may lead to such values on the individual level.

### Fitting the Model (Two-parameter fits)

Given our previous results in which flow parsing gains were vastly different between when object and observer motion were in the same direction and when they were in opposite directions (Jörges and Harris 2022), we fitted a second model in which we fitted different flow parsing gains for the Same Direction and Opposite Directions conditions. This model therefore had two parameters (*k_flow_*_−*parsing*,*opposite*_ and *k_flow_*_−*parsing*,*same*_). Comparing between the one- and two-parameter versions of the model allowed us to address our Supplemental Question 1.

### Fitting the Model (Reduced Dataset)

We fitted both the One-Parameter Model and the Two-Parameter Model to Reduced Dataset A in which we excluded those conditions where participants overwhelmingly misperceived the direction of movement of the sphere (i.e., when the sphere had a speed of 3 m/s in the same direction as the observer). If such an exclusion yielded improved model fits, then that would constitute evidence that our simple flow parsing model captured performance in the excluded condition less well than in the other conditions. This would then indicate that a different mechanism may be at play.

For the sake of comparison, we also fitted the model to partial datasets where we excluded the other combinations of motion profile and sphere speed (Reduced Dataset B: excluding Same Direction, 10 m/s; Reduced Dataset C: excluding Opposite Directions, 3 m/s; Reduced Dataset D, excluding 10 m/s). This helped us answer Supplemental Question 2.

### Modelling Results

When fitting the model to the whole data set, we obtained model fits of R^2 = 0.76 for the Two- Parameter model and R^2 = 0.72 for the One-Parameter model. Model fits were substantially better than this when the two models were fitted to Reduced Dataset A (0.93 and 0.82 for the one- and two-parameter fits respectively) Reduced Dataset B (0.92 and 0.86 for the one- and two-parameter fits respectively). For the Reduced Datasets C and D, model fits decreased or stayed the same (C: 0.73 and 0.68, D: 0.74 and 0.72 for the one- and two-parameter fits respectively). The model fits (in this case, model predictions in comparison to the data) for reduced datasets A and B are shown in Figure 5.

**Figure 5.**
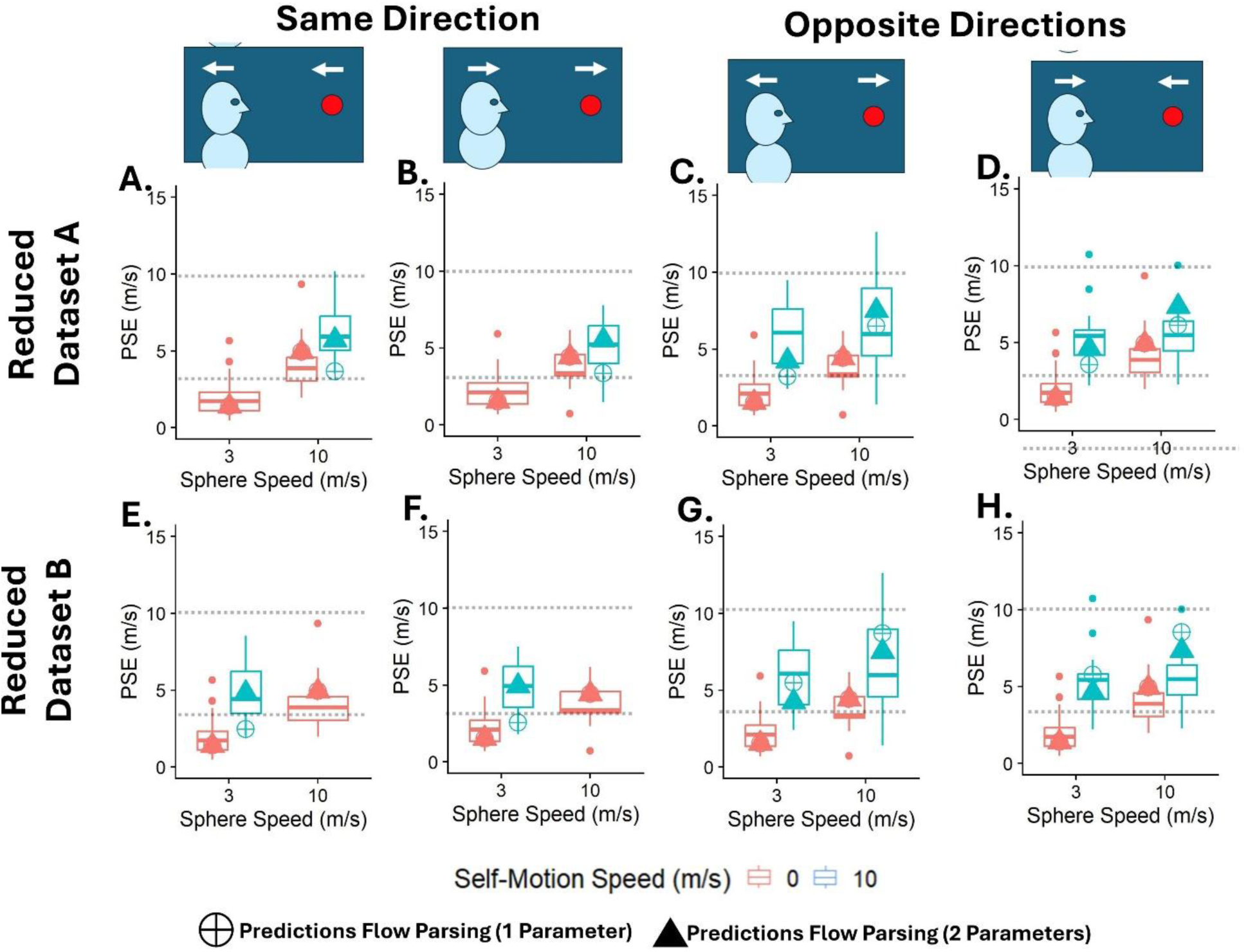
Boxplots of the PSEs for sphere speeds (x axis), separately for all combinations of observer and sphere speeds (panels) and the self-motion speeds (color-coded). The overlayed shapes are the predictions for the One-Parameter flow parsing model (crosshair) and the Two-Parameter flow parsing model (triangle). A & E. For observer and sphere motion out of the scene, i.e., towards the back of the observer. B & F. For observer and sphere motion into the scene, i.e., in the direction the observer was facing. C & G. For backward observer motion and sphere motion into the scene, i.e., in the direction the observer was facing. D & H. For forwards observer motion and sphere motion out of the scene, i.e., towards the back of the observer. A, B, C, D for Reduced Dataset A. E, F, G, H for Reduced Dataset B.

This suggests that our model was able to predict PSEs consistently when the sphere and self-motion were in Opposite Directions but for the Same Direction motion profiles, it could only predict PSEs well for either the 3 m/s or 10 m/s sphere speeds, but not both with the same parameters.

Since we were unable to adjudicate which of the Same Direction conditions, 3m/s or 10m/s, was the odd one out based on model fits alone, we estimated the flow parsing parameters using both Reduced Dataset A (excluding Same Direction 3 m/s) and Reduced Dataset B (excluding Same Direction 10 m/s). We used a *likelihood ratio test* (Wilks 1938) to determine whether the second parameter in the two-parameter Model increased the explained variability significantly for either reduced dataset. This turned out to be the case, both for Reduced Dataset A (p < 0.0001) and Reduced Dataset B (p < 0.0001). The fact that the two-parameter model was significantly better than the one-parameter model shows that flow parsing gains were significantly different for self-motion in the same direction as the target than they were for self-motion in the opposite direction. To further evaluate which condition was the odd one out, we looked at the fitted flow parsing gain parameters separately for Reduced Dataset A and Reduced Dataset B. For the two-parameter model we found a median *k_flow_*_−*parsing*,*opposite*_ = 0.2 and *k_flow_*_−*parsing*,*same*_ = 0.56 when fitting to Reduced Dataset A, and *k_flow_*_−*parsing*,*opposite*_ = −0.08 and *k_flow_*_−*parsing*,*same*_ = 0.2 when fitting to Reduced Dataset B. The full distribution of flow-parsing gains is depicted in Figure 6.

**Figure 6:**
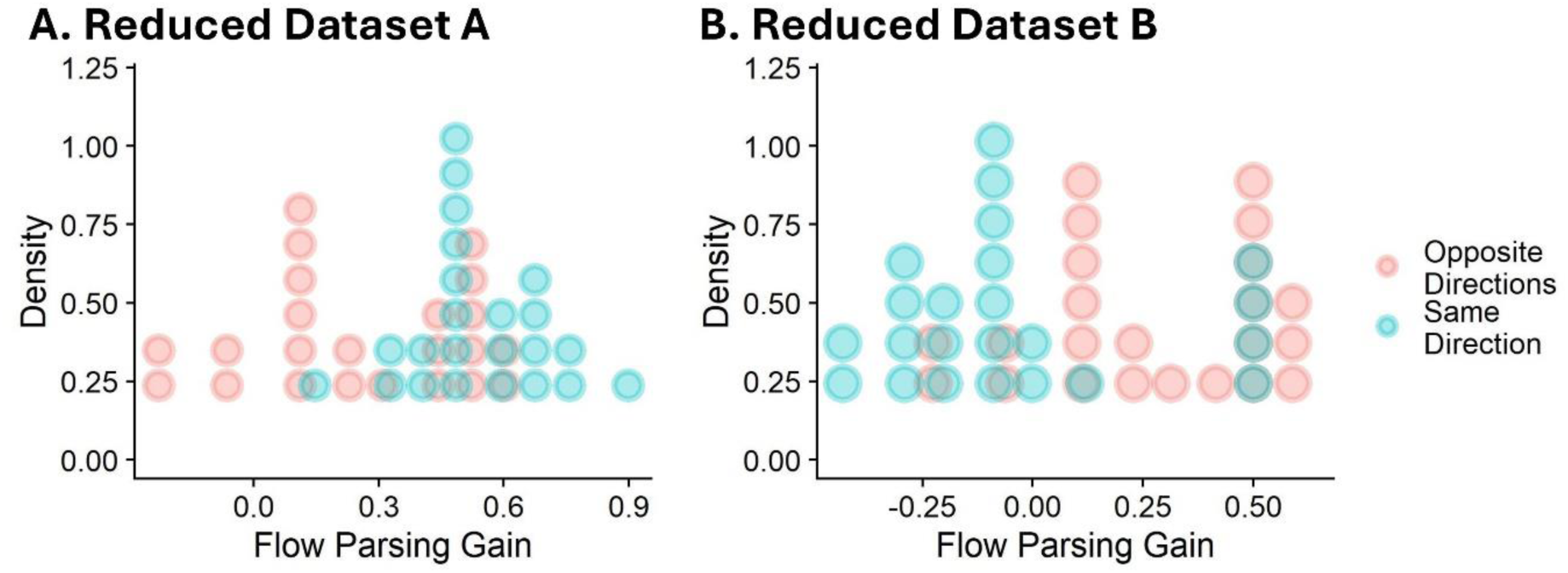
Histograms of flow parsing gains fitted using reduced dataset A (excluding same direction 3 m/s) and using reduced dataset B (excluding same direction 10m/s) using the two-parameter flow parsing model for self-motion in the direction opposite to the target (red) and for self-motion in the same direction as the target (blue). Each dot represents one participant.

### Modelling summary

Overarching Question: A simple flow parsing model can account for the paradoxical finding that object speed was overestimated for the Same Direction motion profile, but only for a sphere speed of 3m/s or a sphere speed of 10m/s, but not for both at the same time. We address the question of which of these two scenarios is likely to be “the odd one out” in the general discussion.

Supplemental Question 1: Our finding that the 2 Parameter model yields significantly better model fits than the 1 Parameter model (even after penalizing for the additional parameter) supports the notion that participants flow parse differently when they move in the same direction as the target than when they move in the opposite direction of the target.

Supplemental Question 2: By showing that our model can explain performance for Same Direction, 3m/s or for Same Direction, 10m/s, but not for both at the same time, we determine that there is a qualitative difference between these two conditions that cannot be captured by a simple flow parsing model.

## Discussion

Curiously, all self-motion conditions were associated with significantly faster perception of ball motion compared to when observers were stationary irrespective of their relative directions. The fact that self-motion in the same direction as the target led to faster PSEs is highly counterintuitive. Self-motion and object motion cancel each other out in this condition, leading to a slower retinal speed and consequently a potential *underestimation* of object speed. An *increased* perceived speed corresponds to an overcompensation for self-motion or more-than-complete flow parsing. This phenomenon has, to our knowledge, never been reported in the literature.

Variations in the parameter ‘*k_relative speed_*’ are likely related to the well-known phenomenon of compression of space in VR (El Jamiy and Marsh 2019; Petukhov et al. 2024). When VR space is compressed, a general underestimation of speed-in-depth would be expected in comparison to lateral speeds. Information about motion-in-depth (Lee et al. 2019) is noisier than for lateral motion (McKee, 1981), which would perhaps make it more susceptible to influences from the slow-motion prior (Stocker and Simoncelli 2006) resulting in slower perceived speeds.

In line with the Flow Parsing hypothesis and previous results (Gray et al. 2004; Rushton and Warren 2005; Warren and Rushton 2008, 2009b; Niehorster and Li 2017; Rushton et al. 2018; Layton and Niehorster 2019; Jörges and Harris 2022, 2024), we predicted that visually simulated self-motion in the same direction as the target would lead to an underestimation of object speed and self-motion in the opposite direction of the target would lead to an overestimation of object speed due to incomplete parsing and the contamination of object speed with the visual consequences of self-motion. While we found strong support for the latter in both experiments, we – perplexingly – found that object speed was overestimated independent of whether self-motion was in the *same* or opposite direction to the object. These paradoxical findings were particularly pronounced for the lower sphere speed of 3 m/s participants were likely to misjudge the direction of the sphere.

We then used a simple geometric model to test whether this paradoxical finding could be a result of differential rates of splitting of optic flow into object speed and self-motion speed estimates. While the model held up well for the opposite-directions motion pairings, for the same-direction motion pairings, it could either account for the same direction 3 m/s condition or the same direction 10 m/s condition but not for both with the same parameters.

While flow parsing has been studied extensively, our present study is one of few that use perceived speed in the world as the variable of interest. To our knowledge, only Hogendoorn et al. (Hogendoorn et al. 2017a) have studied speed judgements. For a rotating observer and 2D motion patches, they reported results predicted by the flow parsing hypothesis in which self-motion was underestimated: that is they reported an overestimation of speed when stimulus motion was opposite to observer motion and an underestimation of speed when stimulus motion was in the same direction as the observer. This symmetry using rotatory motion is contrary to both the present results and our previous results on lateral object and observer motion (Jörges and Harris 2022, 2024). However, the discrepancy may be explained by the fact that Hogendoorn et al. (2017) used physical rotatory motion, whereas our study used visually simulated linear self-motion. When it comes to precision, our results are also in conflict with the Hogendoorn et al. (2017) study: while we found higher JNDs in the presence of self-motion as predicted by the flow parsing hypothesis, Hogendoorn et al. (2017) were unable to detect any impact of self-motion on JNDs. Once again, the reason might be that self-motion was experienced multimodally by the Hogendoorn et al. (2017) participants, while in our case self-motion was simulated only visually. The additional information provided by the vestibular system may have compensated for any precision penalties incurred in the presence of self-motion, thus rendering the effect undetectable.

The Flow Parsing gains (20% for opposite directions and 56% for same direction) found in our study appear to be very low in comparison to the literature, with values of 47% (visual-only) and 58% (visuo-vestibular presentation) reported by Dokka et al. (2015), 50-100% reported by Warren & Rushton (2009b), 35% (under impoverished visual conditions) and 60-65% (with more complete visual information) reported by Niehorster and Li (2017), and 70% (opposite directions) to 100% (same direction) reported by Jörges & Harris (2024). This is particularly surprising as our display was richer and more realistic than the majority of displays used in earlier studies, which often used more abstract visual scenes like radial flow displays Dokka et al., 2015; MacNeilage et al., 2012; Rushton & Warren, 2005; Warren & Rushton, 2008a, 2009a). Even when restricting the comparison to studies on self-motion in depth, which could be considered more biologically relevant but also computationally more challenging than, e.g., lateral or rotational motion, we find notably lower flow parsing gains (see, e.g., Niehorster & Li, 2017 who found gains of 60-65%). Despite our more realistic stimulus, it was hard for our participants to accurately estimate their self-motion speed from the optic flow they were experiencing. This may reflect a difference between the present paradigm and earlier flow-parsing studies of self-motion in depth, in which self-motion in depth was typically paired with vertical object motion (Warren and Rushton 2009b; Niehorster and Li 2017) rather than with object motion along the same depth axis as the observer’s movement. While the absolute values are vastly different, the discrepancy between self-motion in the opposite direction of the target and self-motion in the same direction as the target are reminiscent of our earlier studies (Jörges and Harris 2022, 2024), where we also found flow parsing to be more complete for same direction self-motion than for opposite directions self-motion.

The most perplexing feature of the present dataset is the unexpected behavior for the Same Direction motion profile: rather than decreasing perceived speed, as expected under the flow parsing hypothesis, PSEs were increased for both sphere speeds. A reversal of the perceived direction for the slower sphere speed could partially explain this phenomenon; however, our model comparisons showed that flow parsing gains that produced predictions matching performance for the 3 m/s spheres threw model fits off for 10 m/s and vice-versa, while trying to fit both at the same time made for subpar fits for both conditions. Which of these two conditions is, then, the odd one out? *A priori*, both conditions differ in different ways from the remaining conditions (No Self-Motion and Opposite Directions): for spheres moving at 3 m/s in the same direction as the simulated self-motion, participants “caught up” to the sphere over the course of a trial, with a relative speed that shortened the distance between object and observer. This speed differential led a large majority of the participants to misjudge the direction of the object. For the 10 m/s self-motion condition, on the other hand, observer and sphere moved such that the object did not change its position on the retina at all, thus allowing participants to essentially ignore the object and solve the task by merely referring to their self-motion speed. *Empirically*, the model fits were roughly matched when either of the two problematic conditions were excluded. However, flow parsing gains obtained by fitting the model after excluding the Same Direction, 10 m/s condition, were still out of step with the reports in the literature; the negative gains found for the Same Direction motion profile are furthermore theoretically dubious, as it would indicate that participants either misjudged the direction of their self-motion during these trials, or that they subtracted a fraction of their perceived self-motion rather than adding it to the retinal patterns caused by the moving sphere. For these reasons, it is our opinion that a different mechanism might be at play in the Same Direction, 3 m/s condition than for the remainder of the dataset. Future research could address this open question, for example by using a more fine-grained range of self-motion and object motion speeds. Further, it would be useful to improve the signal-to-noise ratio by making the task easier, as perceptuo-decisional difficulties in comparing speed-in-depth to lateral speed may have led participants to fall back on regression to the mean-based estimates (as observed for diverse perceptual processes, e.g., Olkkonen et al. 2014; Aston et al. 2021; Johnen and Zimmermann 2025). Finally, future research could further expand on our second experiment by also including independent judgements of self-motion speed and/or direction.

## Acknowledgements

BJ was funded by a Postdoctoral Transition Fellowship awarded by the Center for Vision Research and the VISTA project at York University, Toronto. LRH was supported by a grant from the Canadian Natural Sciences and Engineering Research Council. AN was supported by the York University Biology Graduate Program and a Connected Minds Master’s Student Scholarship.

## Open Science

All materials, data, and code for analysis, as well as a video of the stimulus can be found in this Open Science Foundation repository: https://osf.io/ty7cg/

